# “Green pointillism”: detecting the within-population variability of budburst in temperate deciduous trees with phenological cameras

**DOI:** 10.1101/771477

**Authors:** Nicolas Delpierre, Kamel Soudani, Daniel Berveiller, Eric Dufrêne, Gabriel Hmimina, Gaëlle Vincent

## Abstract

- Phenological cameras have been used over a decade for identifying plant phenological markers (budburst, leaf senescence) and more generally the greenness dynamics of forest canopies. The analysis is usually carried out over the full camera field of view, with no particular analysis of the variability of phenological markers among trees.
- Here we show that images produced by phenological cameras can be used to quantify the within-population variability of budburst (WPVbb) in temperate deciduous forests. Using 7 site-years of image analyses, we report a strong correlation (r²=0.97) between the WPVbb determined with a phenological camera and its quantification through ground observation.
- We show that WPVbb varies strongly (by a factor of 4) from year to year in a given population, and that those variations are linked with temperature conditions during the budburst period, with colder springs associated to a higher differentiation of budburst (higher WPVbb) among trees.
- Deploying our approach at the continental scale, i.e. throughout phenological cameras networks, would improve the understanding of the spatial (across populations) and temporal (across years) variations of WPVbb, which have strong implications on forest functioning, tree fitness and phenological modelling.

## Introduction

In the temperate and boreal climate zones, the flushing-out of leaves from dormant buds in spring (*alias* ‘budburst’) is a key step in the seasonal cycle of trees’ activity. It marks the start of the carbon acquisition and water loss (photosynthetic) period and is in close relation with the tree’s other organs or tissues seasonal growth and resource acquisition (reviewed in Delpierre *et al.*, 2016b). To this respect, budburst is hypothesized to influence tree growth (to what extent is not clear, see e.g. Čufar *et al.*, 2015; Bontemps *et al.*, 2017; Delpierre *et al.*, 2017). In some temperate angiosperms (e.g. deciduous oaks), the timing of flowering, which is closely related to the flushing-out of leaf buds (Franjic *et al.*, 2011), influences the production of fruits (Lebourgeois *et al.*, 2018; Schermer *et al.*, 2019). This makes budburst an essential trait for the tree functioning, a trait subject to natural selection (Ducousso *et al.*, 1996; Savolainen *et al.*, 2007). Yet, budburst is a very variable trait in forest tree populations. The duration of the budburst period (from the first to the last tree leafing-out in a given year) in temperate forest tree populations is about three weeks (19 days, averaged over 67 populations of *Quercus petraea* (Matt.) Liebl., *Quercus robur* L. and *Fagus sylvatica* L. in Europe; Delpierre *et al.*, 2017). This is about 30% of the amplitude of the continental gradient of budburst observed for those species (Delpierre *et al.*, 2017). The high within-population variability of budburst (WPVbb) in natural tree populations is probably related to biotic (herbivores and pathogens) and abiotic (frost) fluctuating selection pressures that contribute to maintaining high genetic variation on this trait (Crawley & Akhteruzzaman, 1988; Alberto *et al.*, 2011; Dantec *et al.*, 2015).

Interestingly, WPVbb varies across tree populations (Salmela *et al.*, 2013), and across years for a given population (Denéchère *et al.*, 2019). The within-population standard deviation of budburst averaged 4.0 days, but ranged from 1.7 to 9.7 days in 14 populations of nine temperate deciduous tree species (Denéchère *et al.*, 2019). This variability was related to temperature conditions during the budburst period, with colder conditions associated to an increased WPVbb (Denéchère *et al.*, 2019). Whatever their causes, the year-to-year variations of WPVbb have potentially strong ecological implications, influencing the competition of trees for resource (light, water and nutrient) acquisition and transfer throughout the food web (van Dongen *et al.*, 1997; Thackeray *et al.*, 2016). Unfortunately, WPVbb has seldom been documented to date in natural tree populations (but see Denéchère *et al.*, 2019), probably because its quantification remains laborious based on ground phenological observations, which are still needed for observing individual trees (Chesnoiu *et al.*, 2009; Cole & Sheldon, 2017; Delpierre *et al.*, 2017). Indeed, quantifying WPVbb requires observing bud development on a relatively high number of trees per population (*ca*. 30, see Denéchère *et al.*, 2019), which is rarely attained in most phenological studies. Beside this sampling requirement, it is also more demanding in terms of the number of observations campaigns during spring, since one has to “wait” for all trees to burst buds, whereas classical phenological studies typically record the date at which 50% trees of the surveyed population have reached budburst.

Phenological cameras (hereafter phenocams) have been used for over a decade to monitor the phenology of forest canopies (Richardson *et al.*, 2007; Ahrends *et al.*, 2008; Richardson, 2019). They are a very appealing, automated alternative to ground phenological observations. Basically, phenocams periodically (e.g. every hour) take a RGB picture of the canopy. The pictures are post-processed (Filippa *et al.*, 2016) to extract colour indices quantifying continuously the “colour-state” of the canopy, from which particular phenological metrics (e.g. budburst or leaf senescence) can be inferred. The comparison of budburst dates obtained from ground observations and from phenocams are usually good (e.g. Keenan *et al.*, 2014; Xie *et al.*, 2018). To date, the potential of phenocams has mostly been assessed at the canopy scale, corresponding to the camera field of view (Keenan *et al.*, 2014; Klosterman *et al.*, 2014). Some studies have also considered the scale of individual trees (Ahrends *et al.*, 2008; Berra *et al.*, 2016; Kosmala *et al.*, 2016; Xie *et al.*, 2018), but those studies pointed to tree inter-specific differences, not pointing particularly to the within-population (i.e. intra-specific) variability of phenology and the characterization of its inter-annual variability. Here, we explore the potential of phenocams to investigate WPVbb, targeting two research questions: (1) can phenological cameras be used to quantify the year-to-year variations of WPVbb in deciduous forest tree populations?, (2) do WPVbb determined from phenocams show a similar temperature response to the one established previously from ground observations?

## Material and methods

### Study sites and phenological ground observations

We monitored the development of buds in two sessile oak (*Quercus petraea* (Matt.) Liebl.) populations located in the state-owned forests of Barbeau and Orsay, 50 km from each other in the South of the Paris area, France (Table 1). In both populations, we monitored budburst for more than 15 years following an ‘extensive’ sampling. According to this sampling, we monitored bud development over a large number (>100) of individual dominant oaks, from early signs of budburst (typically in mid-March) until 50% of them have reached budburst (yielding the median date of budburst at the population scale). We considered that a tree had reached budburst when 50% of its crown showed open buds. A bud was considered open when the limb of one out of the *ca*. 12 leaves preformed in the bud (Fontaine *et al.*, 1999) was unfolding as visible from the ground (all observations were made with high-magnification binoculars, minimum x8). In this ‘extensive’ sampling, we did not follow the spread of budburst in crowns of particular trees. Instead, we picked randomly >100 dominant oaks in the forest for each observation campaign. Besides the ‘extensive sampling’, we applied for some years an ‘intensive’ sampling over our two populations, focusing on 27 to 66 tagged, dominant oaks (depending on the site and observation year, Table 1) for which we followed the spread of budburst (from all dormant buds to 100% open buds) in each tree crown, typically from mid-to early-May. The ‘extensive’ sampling yielded the population median date of budburst for each oak population over the whole study period (2012-2018). For the years when we applied the ‘intensive’ sampling, we could compute both the median budburst date and the among-trees standard deviation of the budburst date (SD_ground_, expressed in days). The standard deviation is a measure of the average duration between each tree individual budburst date and the average date established over all individuals. In the following, we consider SD as our metric for quantifying WPVbb. For both the ‘extensive’ and ‘intensive’ samplings, we conducted our observations from once a week during the dormant phase to three times a week during the actual budburst period (average time resolution was 3.8 days in Barbeau and 2.5 days in Orsay).

**Table 1.**
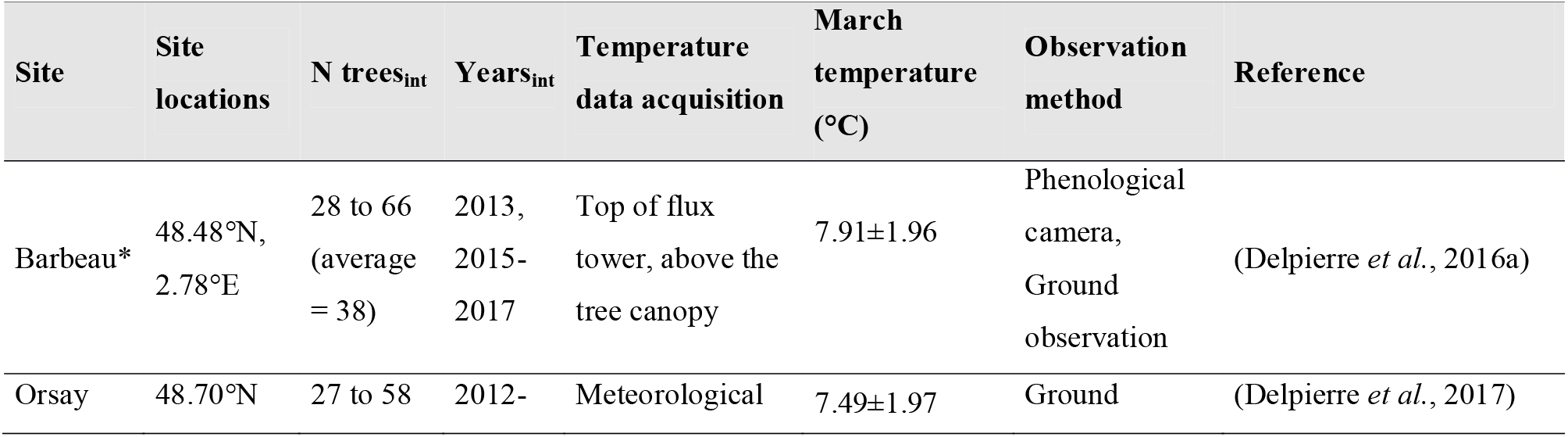
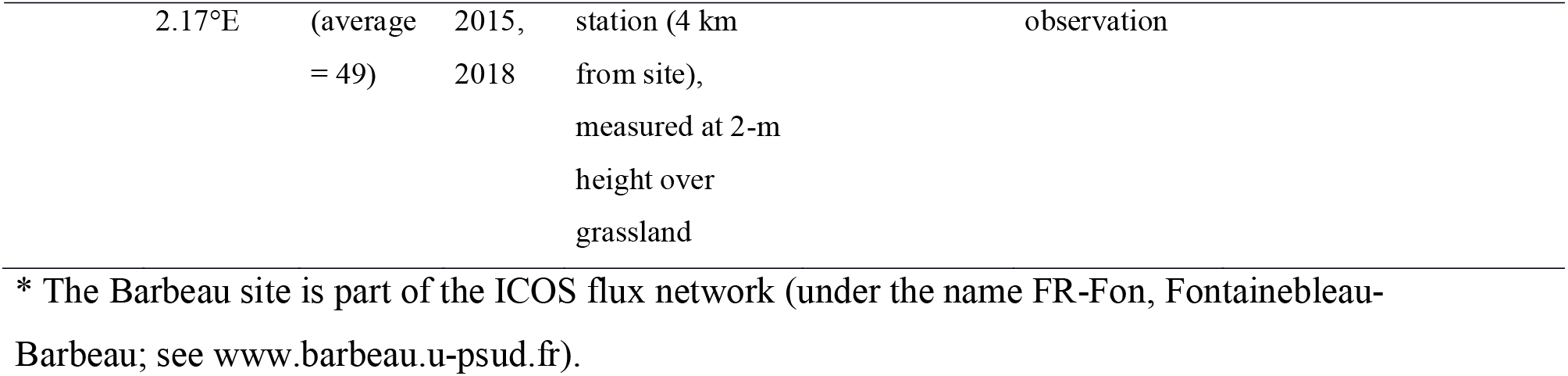
Characteristics of the study sites. *N trees*_*int*_ and *Years*_*int*_ report the number of individual trees and the years for which we applied the ‘intensive’ phenological sampling at a particular site (see text for details). The ‘extensive’ phenological sampling was applied at both sites over the whole study period (2012-2018). At Barbeau, the phenological camera was run over 2012-2018. March temperatures are averages ± SD for 2012-2018.

### Phenological camera and RGB data processing

In the Barbeau forest, we installed a phenological camera (model P1347, Axis Communications, Lund, Sweden) on April 4 (DoY 95), 2012 which has been running continuously since then, covering the whole study period (2012-2018). The camera is mounted at the top of the Fontainebleau-Barbeau flux tower (see Delpierre *et al.*, 2016a for site details), five meters above the tallest trees. RGB pictures of the forest canopy (resolution of 2590 x 1920 Px), were acquired continuously every hour from 8 am to 5 pm (UT), yielding 10 images per day from year 2013. In year 2012, only 1 image per day (at 10 am) was recorded. In order to access to the within-population variability of budburst, thirty regions of interest (ROIs; 43 kPx on average, range 16-102 kPx) were delineated among 16 visible tree crowns from the top of the canopy on a spring image (Fig. 1). In order to minimize effects of changing illumination conditions, two small ROIs were delineated on a white PVC sheet installed in the camera field of view and used as a white reference standard (3 kPx and 1.4 kPx, respectively; Fig. 1). To convert radiance to pseudo-reflectance, the Red, Green and Blue radiance averages of each ROI were respectively divided by the R, G and B radiance averages of the two white standards. These pseudo-reflectances (*ρ*) were averaged on a daily basis (10 values per day, corresponding to the hourly sampling) and used to determine a daily greenness index for each ROI, as: *Gi = ρG*/(*ρR + ρG + ρB*).

**Figure 1.**
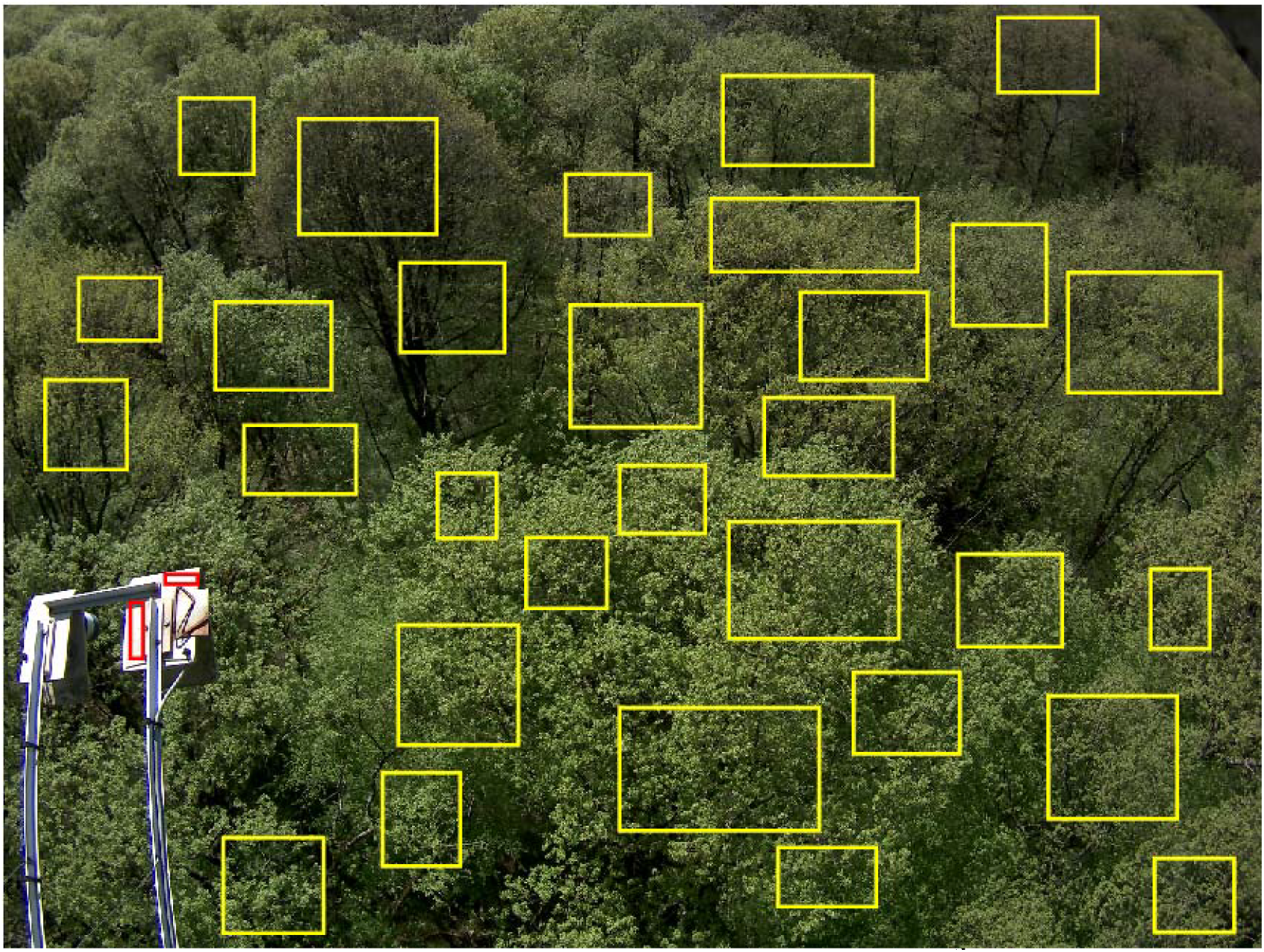
View from the Barbeau phenological camera, on April 19^th^ 2018 (4 pm). The camera is mounted on the platform of the FR-Fon flux tower, 5 meters above the tallest oaks. Yellow rectangles mark the position of 30 regions of interest (ROIs). Red rectangles identify the white reference standards used to calculate RGB pseudo-reflectances. See text for details.

### Extraction of RGB-based phenological metrics

For each ROI and each year, we extracted the date of spring transition from a sigmoid curve fit to Gi time-series (Soudani *et al.*, 2008). The sigmoid curve equation is:

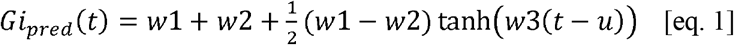

where *Gi*_*pred*_*(t)* is the predicted greenness index at day of year *t*; *tanh* is the hyperbolic tangent and *w1*, *w2*, *w3*, and *u* are the fitting parameters. (*w1*+*w2*) is the minimum *Gi*_*pred*_, reached in the non-leafy season. (*w1-w2*) is the total amplitude of *Gi*_*pred*_ temporal variations. Parameter *u* is the date (DoY) corresponding to the highest rates of change of *Gi*_*pred*_(*t*) (maximum peak of the first derivative of *Gi*_*pred*_(*t*), i.e. the inflexion point, corresponding to 50% of the spring greenness amplitude). There is no consensus in the literature as regards the most appropriate way to quantify budburst from Gi time series. Here, we considered *u* as our proxy for the budburst inferred from phenocam images. Since we were interested in the spring phase, we considered Gi series acquired from DoY 1 to DoY 240. We fitted eq. 1 by minimizing the sum of squares of differences between Gi_pred_ and the measured Gi values using MATLAB v8.5. Fitting eq. 1 to each ROI for each year, we ended up with 30 (ROIs) times 7 (from 2012 to 2018) estimates of budburst dates inferred from the phenocam.

## Results

The population median budburst dates determined according to the ‘extensive’ sampling averaged DoY 104.3 in Barbeau and DoY 104.7 in Orsay over 2012-2018 (Table 2). The dates observed at both populations varied from DoY 97 to 114 and differed at maximum by two days for a particular year. The distributions of the Barbeau and Orsay median budburst dates determined according to the extensive sampling were virtually equal (Student t=-0.13, p<0.89). The ‘intensive’ sampling yielded median budburst dates within one-day of those determined by the ‘extensive’ sampling, at both populations (Table 2). Overall, these results show that both the Barbeau and Orsay populations show virtually identical budburst dates, whatever the sampling scheme considered. At Barbeau, we could determine WPVbb from ground phenological observations for 4 out of 7 years from 2012 to 2018 (Table 1), hence missing 3 years. Considering the high similarity of budburst dates observed for the Orsay and Barbeau site, we proposed to use data from the Orsay ‘intensive’ sampling campaigns as a proxy for those missing at Barbeau (Table 2) for comparison with the phenocam dates.

**Table 2.** Budburst dates of sessile Oak over the study sites. These are the median dates (in day of year, DoY) of budburst determined over the Barbeau and Orsay populations, located 50 km apart. The median date has been determined with two sampling schemes (see text for details). Dash denotes missing data.

| Year | Barbeau<sub>extensive</sub> | Barbeau<sub>intensive</sub> | Orsay<sub>extensive</sub> | Orsay<sub>intensive</sub> |
| --- | --- | --- | --- | --- |
| 2012 | 101 | - | 103 | 103 |
| 2013 | 114 | 114 | 114 | 113 |
| 2014 | 98 | - | 97 | 98 |
| 2015 | 105 | 105 | 105 | 104.5 |
| 2016 | 109 | 108 | 110 | - |
| 2017 | 98 | 98 | 97 | - |
| 2018 | 105 | - | 107 | 108 |

The dynamics of the greenness index in ROIs (Fig. 2a, b) and the percentage of open buds in individual oak crowns (Fig. 2c, d) were similar, both for year of low (2015) or high (2017) WPVbb. At Barbeau, the median budburst date observed from the ground was close to the estimate from the phenocam data processing (RMSD= 4.8 days, n=7; reduced to 1.2 days when excluding year 2012, n=6; Fig. 3a), the latter averaging DoY 105.7 over 2012-2018.

**Figure 2.**
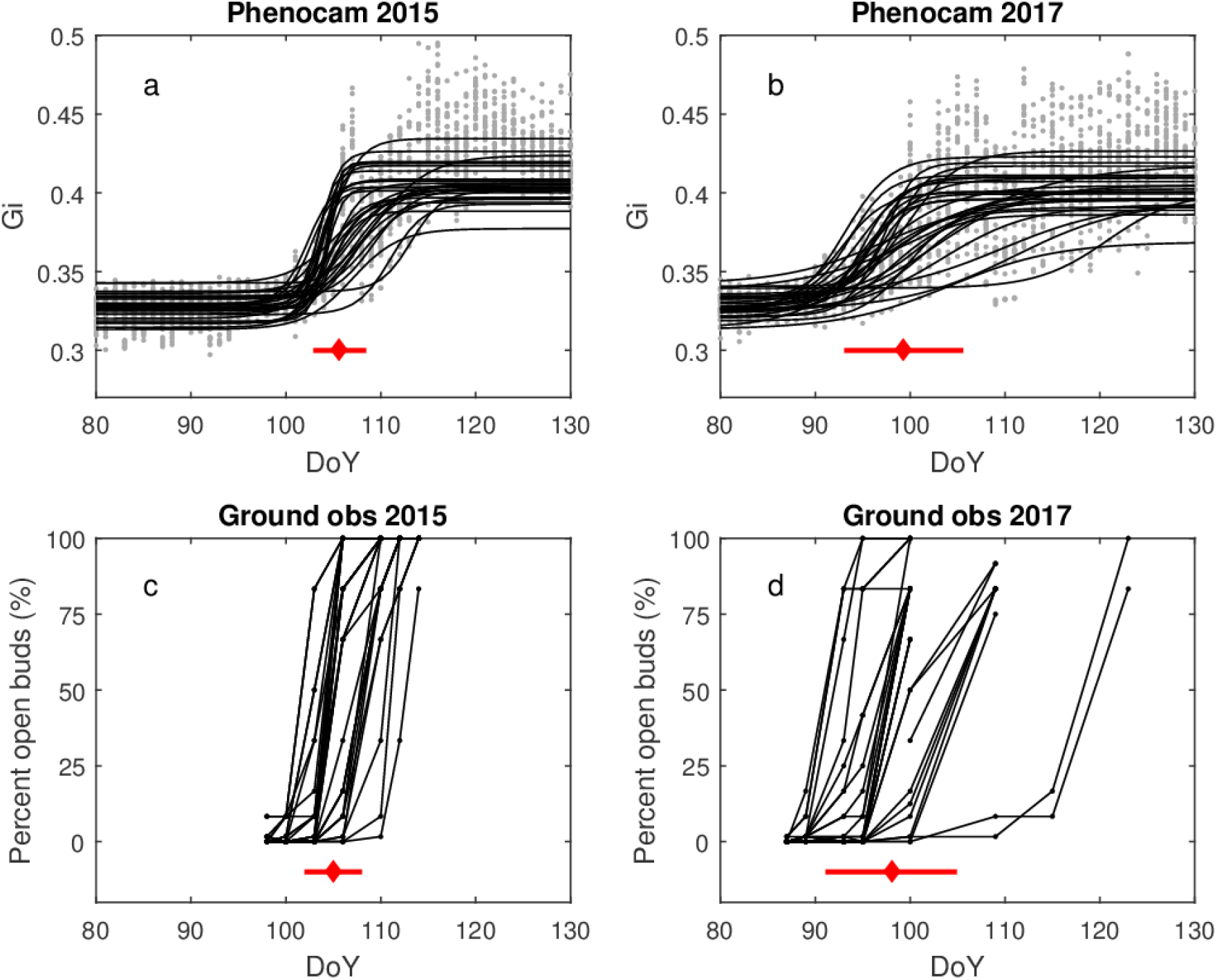
Comparing the dynamics of Greenness index (a, b) and percent open buds observed from ground (c, d) of oaks at the Barbeau forest. Data are from years 2015 (a, c) and 2017 (b, d) which are representative of a year with a low (2015) and a high (2017) WPVbb. On plots (a) and (b), grey dots are the actual data (30 ROIs) and black lines are fits from eq. 1 (one curve per ROI). On plots (c) and (d), black dots are individual observations of percent open buds in tree crowns, with data from one tree joined by a continuous line (n=31 individual trees in 2015, n=30 in 2017). On each subplot, the red diamond marks the median date of budburst determined across ROIs (a, b) or individual tree crowns from the ground (c, d). The horizontal red lines crossing the diamond represent plus/minus one standard deviation. The position of red symbols on the ordinate axis is arbitrary.

**Figure 3.**
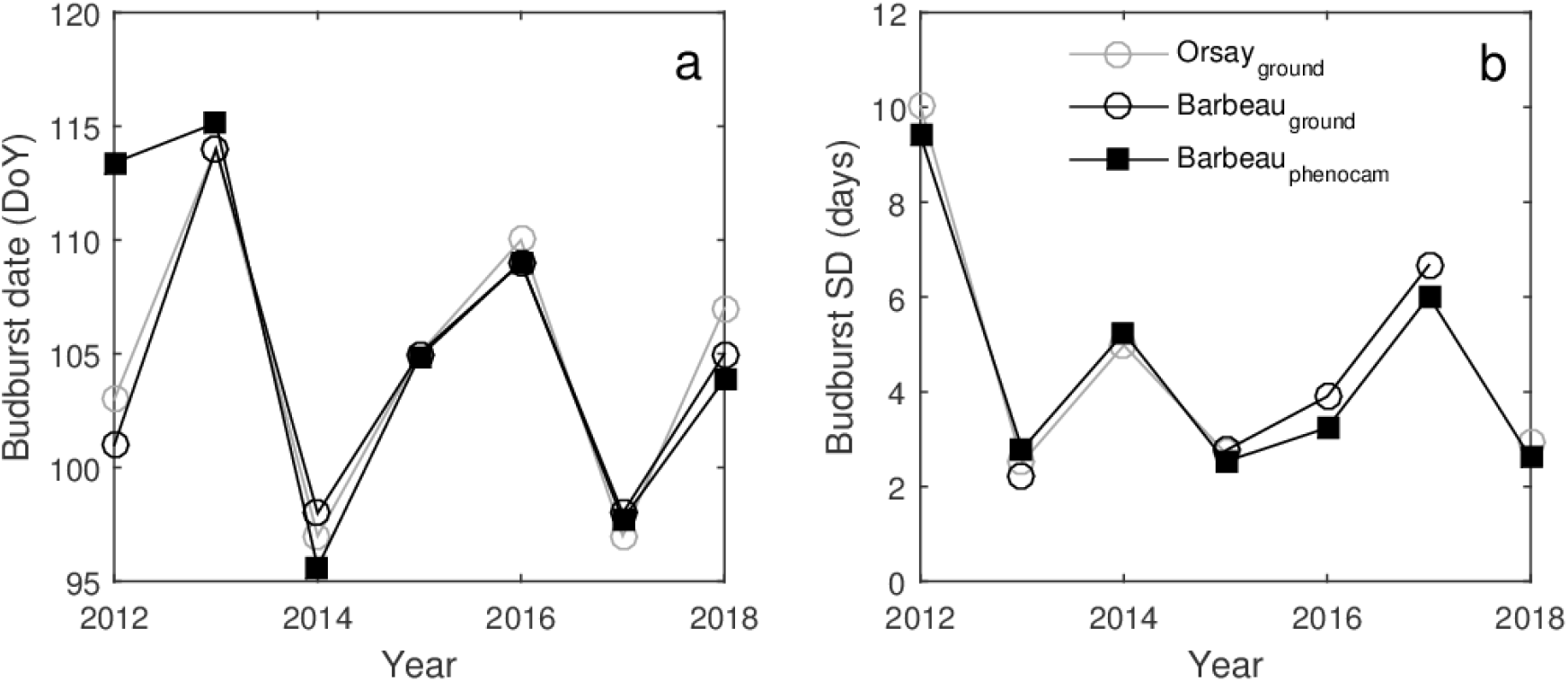
Comparing the population median budburst date (a) and standard deviation (b) for the Orsay and Barbeau populations. The ground observation data shown in (a) were acquired according to the ‘extensive’ sampling scheme.

The standard deviation of budburst calculated from the Barbeau phenocam (SD_cam_) ranged from 2.5 days in 2015 to 9.4 days in 2012, and averaged 4.5 days over 2012-2018 (Fig. 3b). The SD_cam_ values were close to the estimates established from Barbeau ground observations (RMSD= 0.56 days, n=4, Fig. 3b). The standard deviations of budburst observed from the ground in Barbeau and Orsay compared well (RMSD= 0.21 days, n=2) (Fig. 3b). SD_cam_ was strongly correlated, and mostly unbiased, with the SD series obtained from the ground in Orsay and Barbeau (Fig. 3b and Fig. 4).

**Figure 4.**
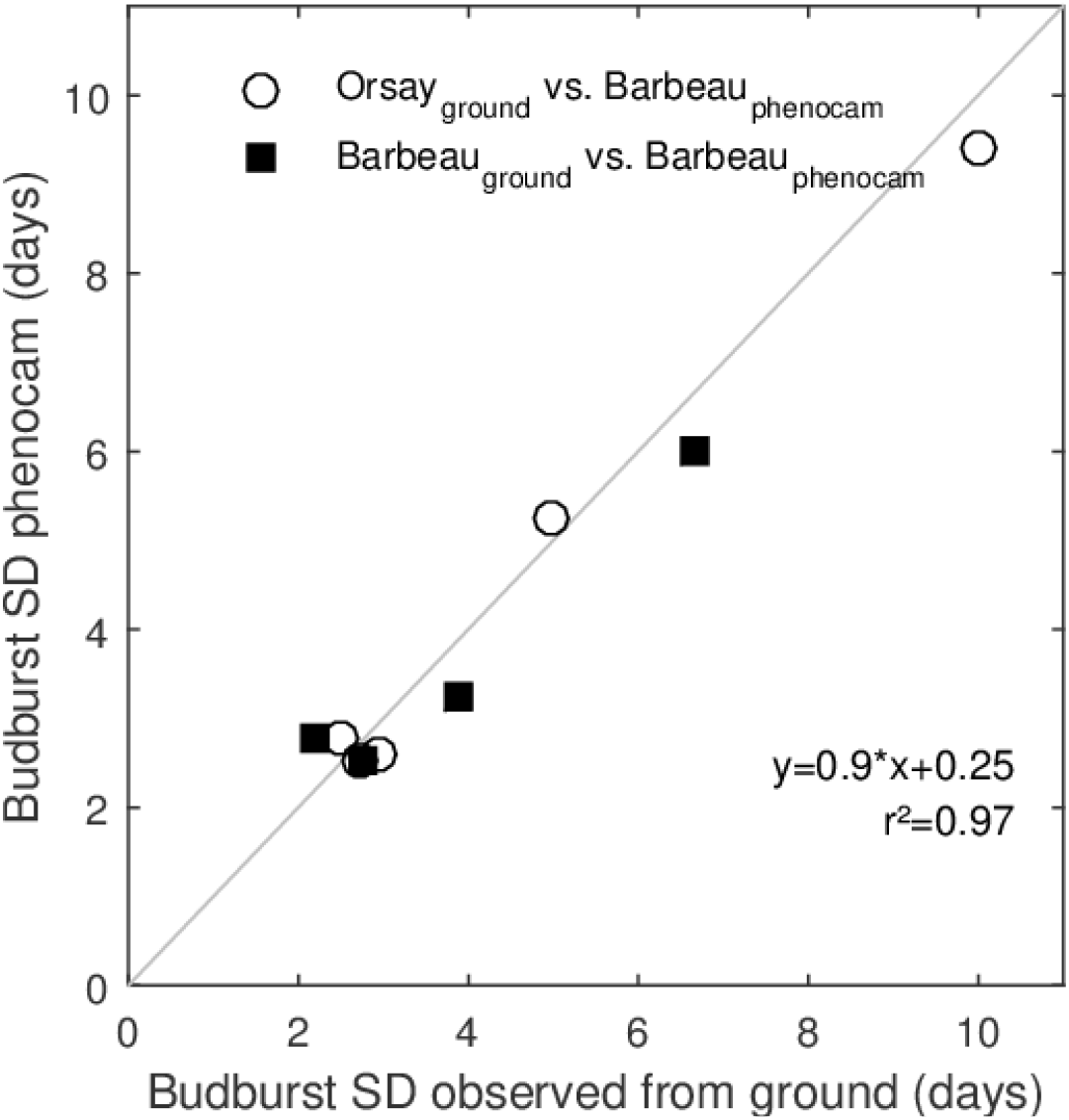
The standard deviation of budburst estimated from the phenocam data processing align with those observed from the ground. Individual points represent one site-year. The linear equation displayed on the graph is established over all data points. The identity line appears in grey.

Either determined from ground observations (in Orsay and Barbeau) of from processing phenocam images (in Barbeau), the standard deviation of budburst was negatively related with the minimum temperature occurring during the budburst period (defined as the time from the first to the last tree bursting its buds in the population sample) (Fig. 5; Spearman’s rank correlation coefficient ρ=-0.66; p<0.006).

**Figure 5.**
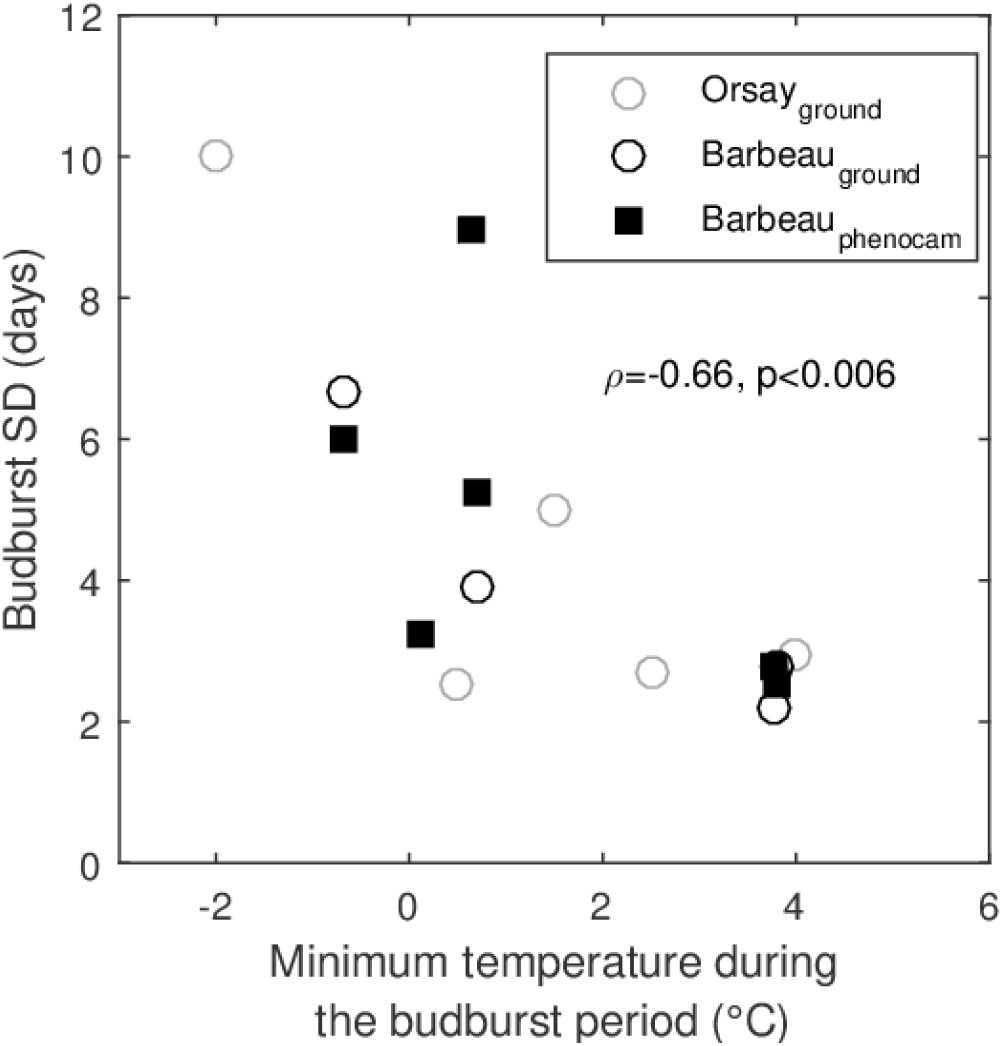
The standard deviation of budburst is related to minimum temperatures during the budburst period. Individual points represent one site-year.

## Discussion

Phenological cameras have been used multiple times to detect phenological transitions from tree individuals to the landscape scale, with errors in the identification of budburst ranging from 1.7 to 9 days (Table 3). Those differences between ground observations and phenocam-derived estimates average 4.5 days (Table 3), and are comparable to the time resolution of ground phenological observations (usually once to three times a week in spring). Here we observed a 4.8 days root mean squared difference between the date of budburst determined with phenocam and the one observed from the ground over of the 7-year time series at the Barbeau forest (Fig. 3a). We notice a lesser comparability of the ground observations and phenocam-derived budburst date for year 2012 at Barbeau (phenocam lags 12 days behind ground observations; Fig. 3a). This was the year when we started phenocam data acquisition, and we installed the camera relatively late (on DoY 95, see Material and Methods), after the first trees has leafed-out in our population. In fact, data from our ‘extensive’ sampling at Barbeau indicate that on this very date, 38% of 210 trees had already burst their buds (data not shown).

**Table 3.** Survey of published literature reporting comparisons between ground phenological observations and spring phenological transition determined with phenocams. Gcc= green chromatic coordinate; FOV= field of view.

| Reference | RMSD (days) | N | Phenocam method | Independent observation method |
| --- | --- | --- | --- | --- |
| (Ahrends *et al.*, 2009) | 3-4 | 7 ROIs in the same FOV, 2 years | Gcc spring, 50% greenness amplitude | Visual assessment of phenocam images (looking for “visible leaf-out”) |
| (Keenan *et al.*, 2014) | 1.7 | 5 years | Gcc thresholding (=0.38) | Ground observation of budburst (95% open buds) averaged over 4 individuals |
| (Klosterman *et al.*, 2014) | 7-9 | 73 site-years | Various sigmoid models fitted to Gcc, 10% greenness amplitude | Visual assessment of phenocam images (majority of trees start to leaf-out) |
| (Klosterman & Richardson, 2017) | 4.7 | 30 tree individuals (1 year) | Sigmoid model fitted to Gcc, 10% greenness amplitude | Ground observations of 10% budburst for 30 trees |
| This study | 4.1 (1.1) | 7 years | Sigmoid model fitted to Gi, 50% greenness amplitude | Ground observations of 50% budburst for >100 trees |

There is no consensus in the literature as regards the way to process phenocam images to detect the date of budburst (Table 3). In this study, we used the inflection point of a sigmoid model fitted to the spring Gi time series as our phenocam-derived metric of budburst. Our results show that this method compares well to the median date of budburst observed from the ground over the Barbeau oak population (Fig. 3a). More generally, we notice that there is no universal protocol for the ground observation of phenological transitions at the scale of tree crowns. The BBCH scale (Finn *et al.*, 2007), often considered as a reference for phenological observations is defined at the scale of organs (buds, leaves), and there is no common protocol at higher (i.e. the tree crown) scale. Phenological cameras are appealing as they offer a way to process imagery signals in a uniform way (Richardson *et al.*, 2018), going over uncertainties associated with the acquisition of data by multiple human observers (Cole & Sheldon, 2017); Liu et al. in prep). On the other hand, phenological metrics derived from phenocams need comparisons with ground-truth phenological data, at least for making them comparable with the multi-decade phenological time series acquired in the field (Templ *et al.*, 2018).

Our main objective here was to assess whether the within-population variability of budburst (WPVbb) could be detected with phenocam data. For this purpose, we needed a ground-truth quantification of WPVbb. Quantifying WPVbb with ground phenological observations is time-consuming. Indeed, in order to derive a robust estimate of WPVbb, one needs to observe at least 30 trees (Denéchère *et al.*, 2019), with a typical amplitude of around 20 days from the first to the latest tree to burst buds (Delpierre *et al.*, 2017); note that we attained a 35 days amplitude in Orsay 2012, corresponding to our maximum standard deviation of budburst, Fig. 3b). This is why we did not monitor ground phenology over the whole period of phenocam data acquisition at Barbeau (2012-2018, i.e. 7 years), located 70-km away from our lab. We completed this time series with data acquired at Orsay, which allowed us to cover the 7-year period of phenocam observation. The population median dates of budburst compare very well (they are within two days, Table 2) in Barbeau and Orsay over the study period, which is not surprising since both populations are separated by 50 km in plain and essentially experiment the same climate conditions (Table 1). What is more interesting is that the WPVbb determined from ground observations was very similar for both sites and compared well with WPVbb derived from phenocam (Fig. 3b). Beyond the validation of the use of phenocam to detect WPVbb, this result implies that our two oak populations show very similar budburst but also WPVbb. This result is very interesting and mirrors the comparability of oak populations at regional scales (see Suppl. Mat. 1 for a comparison of budburst dates at the continental scale), notably driven by pollen dispersal (Kremer *et al.*, 2002).

The standard deviation of budburst we report here averaged 4.5 days (as derived from the Barbeau phenocam). This is comparable to the average of 4.0 days (range 1.7 to 9.7 days) observed in 14 tree populations of nine temperate deciduous tree species (Denéchère *et al.*, 2019). It is also comparable to the value of 3.5 days detected by the analysis of RGB-derived greenness index time series acquired with an unmanned aerial vehicle (UAV, i.e. a drone) for one year over 60 grid cells (100 m2 each) containing deciduous trees in the Harvard forest (Klosterman *et al.*, 2018). Coniferous tree species may display larger standard deviation of budburst: Salmela *et al.* (2013) report an average value of 7.4 days (ranging from 4.3 to 11.1 days) in 21 populations of Scots pine grown in a common garden. Though being an adaptive trait in Scots pine (Salmela *et al.*, 2013), budburst may undergo less selection pressure for evergreens (that by definition remain at least potentially photosynthetically active throughout winter and spring; Mäkelä *et al.*, 2004) than for deciduous trees.

Our analysis of the Barbeau phenocam data evidenced a high interannual variability of WPVbb for a given tree population (Figure 3b). We observed that lower minimum temperatures during the budburst phase are associated with higher WPVbb (Fig. 5). This result is similar to the one observed across 14 European tree populations (Denéchère *et al.*, 2019). Our hypothesis is that as the accumulation of degree-days occurs faster during a warm spring, the time interval from the first to the last tree bursting buds in the population is reduced as compared to a colder spring (see Denéchère *et al.*, 2019, their suppl. Mat).

## Conclusion and perspectives

Phenological cameras have been used for over a decade to detect spring transition from the dormant to the active phase in temperate deciduous canopies. Here, we demonstrated through comparison with ground observations acquired over 7 years on sessile oak populations that phenological cameras can further be used to quantify and monitor the year-to-year variations of the within-population variability of budburst (WPVbb) in temperate deciduous forests. Generalizing our approach over continental scale phenocam networks (Wingate *et al.*, 2015; Richardson *et al.*, 2018) would increase our understanding of the spatial (i.e. across population) and temporal variability of WPVbb. The implications of considering WPVbb in phenological modelling are two-folds: (1) phenological models are classically built to represent the year-to-year variations of the average date of budburst of a tree population, and we hypothesize that the accuracy of phenological models is lower for years of higher WPVbb; (2) the emerging class of physio-demo-genetic models (Kramer *et al.*, 2008, 2015; Oddou-Muratorio & Davi, 2014), aiming at simulating the micro-evolution of tree populations, needs accurate parameterizations of the within-population variability of leaf phenological traits. Quantifying WPVbb with phenological cameras will help documenting those aspects.

## Acknowledgements

We thank the Groupe d’Intérêt Public sur les Ecosystèmes forestiers (GIP-Ecofor) for continuously supporting research activities at the Barbeau forest. We acknowledge the contribution of temperature data from Météo-France, used for analysing data from the Orsay forest. We thank Félix Roux, Guillaume Douriez and Rémy Denéchère for participating to the ground phenological monitoring. The authors have no conflict of interest to declare.

## Authors’ contribution

ND designed the research and wrote the manuscript. KS processed the phenological camera images. DB and GH operated the phenological camera. ND, ED, GV and DB made ground phenological observations. All the authors commented and approved the paper.

